# Polymorphism of Genetic Ambigrams

**DOI:** 10.1101/2021.02.16.431493

**Authors:** Gytis Dudas, Greg Huber, Michael Wilkinson, David Yllanes

## Abstract

Double synonyms in the genetic code can be used as a tool to test competing hypotheses regarding ambigrammatic narnavirus genomes. Applying the analysis to recent observations of *Culex narnavirus 1* and *Zhejiang mosquito virus 3* ambigrammatic viruses indicates that the open reading frame on the complementary strand of the segment coding for RNA-dependent RNA polymerase does *not* code for a functional protein. *Culex narnavirus 1* has been shown to possess a second segment, also ambigrammatic, termed ‘Robin’. We find a comparable segment for *Zhejiang mosquito virus 3*, a moderately diverged relative of *Culex narnavirus 1*. Our analysis of Robin polymorphisms suggests that its reverse open reading frame also does not code for a functional protein. We make a hypothesis about its role.

## Introduction

Of all the various types of viruses catalogued, narnaviruses rank among the simplest and most surprising [Cobián Güemes et al., 2016]. Narnaviruses (a contraction of ‘naked RNA virus’) are examples of a minimal blueprint for a virus: no capsid, no envelope, no apparent assembly of any kind. The known narnavirus blueprint appeared for all intents and purposes to be a single gene, that which codes for an RNA-dependent RNA polymerase, abbreviated as RdRp, [Hillman and Cai, 2013]. However, some narnaviruses have been found to have a genome with an open reading frame (i.e., a reading frame without stop codons) on the strand complementary to that coding for the RdRp gene, calling into question the general hypothesis of a one-gene blueprint [Cook et al., 2013, DeRisi et al., 2019, Dinan et al., 2020, Cepelewicz, 2020]. This reverse open reading frame (rORF) has codon boundaries aligned with the forward reading frame. Because the genome can be translated in either direction, we say that these narnaviruses are *ambigrammatic*. The significance of an ambigrammatic genome is an open problem. In this paper we discuss how polymorphisms of sampled sequences can distinguish between competing hypotheses on the function and nature of ambigrammatic viral genomes. Our methods are applied to known ambigrammatic narnavirus genes and to the newly discovered ambigrammatic second segment of some narnaviruses, termed *Robin* [Batson et al., 2020].

Our discussion is based upon two rules about the genetic code and its relation to ambigrammatic sequences. Both of these *ambigram rules* are concerned with the availability of synonyms within the genetic code, which allow coding of the same amino acid with a different codon. The first rule states that for any sequence of amino acids coded by the forward strand, it is possible to use individual synonymous substitutions to remove all stop codons on the complementary strand [this result was discussed already in DeRisi et al., 2019]. The second ambigram rule, described below, states that the genetic code contains double synonyms that allow polymorphisms, accessible by single-base mutations, even when the amino acids coded by both the forward and the complementary strands are fixed.

The first of these rules addresses the ‘how’ of ambigrammatic genomes, by showing that stop codons on the complementary strand can be removed by single-point mutations, without altering the protein (in narnaviruses, the RdRp) coded in the forward direction. Here we argue that the second rule can help to resolve the ‘why’ of ambigrammatic genomes: the origin of ambigrammaticity itself. There are two distinct reasons why there might be an evolutionary advantage for a virus to evolve an ambigrammatic sequence. The first possibility is that the complementary strand might code for a functionally significant protein, for example, one that might interfere with host defence mechanisms. The second possibility is that the lack of stop codons on the complementary strand is significant, even if the amino acid sequence that is coded is irrelevant. In particular, the lack of stop codons may promote the association between ribosomes and the complementary strand viral RNA (produced as part of its replication cycle). It is possible that a ‘polysome’ formed by a covering of ribosomes helps to shield the virus from degradation or from detection by cellular defence mechanisms [Cepelewicz, 2020, Retallack et al., 2021, Wilkinson et al., 2021]. The second ambigram rule combined with data on the polymorphism of the virus genome can help distinguish whether the complementary strand codes for a functional protein. We shall argue that in the case of *Culex narnavirus 1* and *Zhejiang mosquito virus 3*, the evidence is in favour of this second hypothesis, namely that the open reading frame (ORF) on the complementary strand does not code for a functional protein.

After describing the genetic ambigram rules, we discuss how the existence of double synonyms can be used to assess whether the open reading frame on the complementary strand codes for functional protein. It is well known that, because RdRp is a highly-conserved gene, CxNV1 non-synonymous mutations are likely to be detrimental, so that most of the observed diversity consists of synonymous changes. Some of these synonymous mutations have the potential to be synonymous in the complementary strand. If the complementary strand also codes for a functional protein, we expect that doubly synonymous mutations will be favoured. In fact, there would be mutational ‘hotspots’ corresponding to the potential doubly-synonymous loci. We introduce two tests for whether the complementary strand is coding, based respectively on looking for mutational ‘hotspots’, and upon the mutational frequencies at loci which have double synonyms. We used these tests to analyse sequences for two different ambigrammatic narnaviruses: 46 RdRp segments of *Culex narnavirus 1* [Göertz et al., 2019] and 12 RdRp segments of *Zhejiang mosquito virus 3* [Shi et al., 2017], abbreviated to CxNV1 and ZJMV3 respectively. We find that neither of our tests supports the hypothesis that the translated sequence of the complementary strand of RdRp is under purifying selection. We also applied these tests to the second segment, termed *Robin*, which is found to be closely associated with this ambigrammatic narnavirus infection in mosquitos [Batson et al., 2020, Retallack et al., 2021]. We also found that the complementary open reading frame of Robin does not appear to be under purifying selection. The discovery of Robin suggested that ambigrammatic companions may exist for other ambigrammatic viruses. Accordingly, we searched the assembled contigs of studies reporting the detection of ZJMV3, the only other ambigrammatic narnavirus observed multiple times in numerous locations, and discovered an ambigrammatic segment with similar properties to CxNV1 Robin. Thus we consider four viral segments, denoted CxNV1-RdRp, CxNV1-Robin, ZJMV3-RdRp, ZJMV3-Robin. We shall report evidence that Robin does code for a protein in its forward direction, but that its complementary strand does not code for a functional protein. We find evidence that Robin segments are under detectable purifying selection. Figure 1 illustrates the phylogenetic relationship of CxNV1 and ZJMV3, and ORF-wide d*N* /d*S* values of all their segments and coding directions (discussed in detail below).

**Fig. 1.**
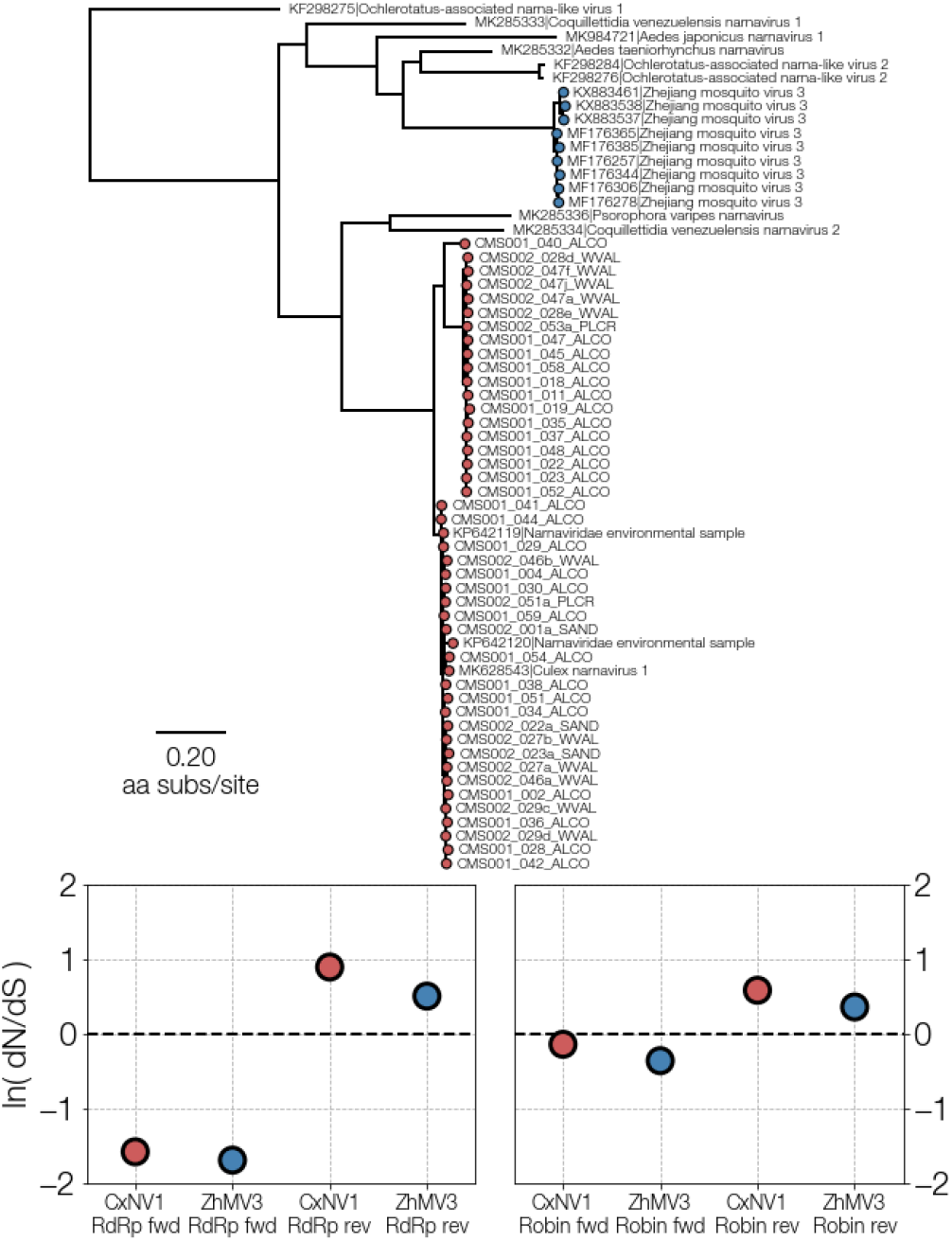
**a** A maximum-likelihood tree illustrating the relationship between CxNV1 (*Culex narnavirus 1*) (red) and ZJMV3 (*Zhejiang mosquito virus 3*) (blue) and related narnavirus RdRp amino acid sequences. **b** ORF-wide d*N* /d*S* values for forward and reverse directions of RdRp and Robin segments for both viruses.

Some careful consideration is required to reconcile our observations with results recently reported in Retallack et al. [2021], where it was shown that introducing mutations which are non-synonymous on the reverse open reading frame of *Culex narnavirus 1* can reduce the fitness of this virus. In the concluding section, we consider the interpretation of these observations, and discuss whether there may be implications for other viral families.

There are many examples of overlapping viral genes with staggered reading frames: this was first clearly described in Barrell et al. [1976], and has been reviewed in Chirico et al. [2010]. Recent work by Nelson, Ardern and Wei [Nelson et al., 2020] discusses how these can be identified. Our investigations indicate that the ambigrammatic ORFs discussed in this work are a different phenomenon, because they do not code for a functional protein. Our approach to analysing the ambigrammatic sequences is quite distinct from the rather complex machinery proposed in Nelson et al. [2020], because it emphasises the role of double synonyms as an unambiguous discriminant of the role of the ambigrammatic sequences.

For completeness, we also include some data on the relative prevalence of the two strands of the CxNV1 virus (both the RdRp and the Robin segments). The results do not appear to be unusual for a narnavirus, which is consistent with the hypothesis that the strand prevalences are not related to the ambigram property.

### Ambigram rules and their significance

We start by describing the two genetic ambigram rules.

**Rule 1** *All complementary-strand stops are removable*

Consider the reading frame on the complementary strand that has its codons aligned with those on the forward strand. Every codon on the forward strand corresponds to a complementary-strand codon read in the reverse direction. The rule states that any stop codon on the complementary strand can be removed by a single-point mutation which leaves the amino acid specified by the forward-read codon unchanged.

This result is demonstrated by the following argument, as discussed in DeRisi et al. [2019]. Reversing the read direction and taking the pairing complement, the stop codons UAA, UAG, UGA in the standard genetic code become, respectively, UUA, CUA, UCA, for which the amino acids are Leu, Leu, Ser. It is only instances of leucine and serine in the forward sequence that can result in stop codons in the reverse read. The synonyms of Leu are CUN, UUA, UUG (where N means any base). The synonyms of Ser are UCN, AGU, AGC. The undesirable Leu codon UUA can be transformed to UUG by a single substitution. Similarly, the Leu codon CUA can be transformed to CUU, CUG or CUC by single substitutions. And the Ser codon UCA is transformed to UCU, UCG or UCC by single substitutions. We conclude that every stop codon on the reverse reading frame can be removed by a synonymous, single site nucleotide mutation.

Furthermore, it is found that complementary-strand stops cannot always be removed by synonymous substitutions in the other two read frames for the complementary strand [each case requires a separate and somewhat involved argument, also given in DeRisi et al., 2019]. As a consequence of these two arguments, we need discuss only the complementary read frame with aligned codons.

**Rule 2** *There exist double synonyms*

Most synonymous mutations of the forward strand produce a non-synonymous change in the complementary strand, but the genetic code does include a number of double synonyms, where the reverse complement of a synonymous mutation is also a synonym. For example codon AGG (Arg) can become CGG (Arg) via a synonymous mutation, while the reverse complement of AGG, which is CCU (Pro) transforms to CCG (Pro) under the same mutation.

The full set of double synonyms in the standard genetic code are as follows:

- Two of the six synonyms of Ser are double synonyms, with reverse complements coding Arg. Conversely, two of the six synonyms of Arg are double synonyms, with reverse complement coding Ser.
- Two more of the six synonyms of Arg are double synonyms, with reverse complement Pro. Conversely, two of the four synonyms of Pro are double synonyms coding for Arg.
- Two of the six synonyms of Leu are double synonyms, with reverse complement Gln. Conversely, both synonyms of Gln are double synonyms, with reverse complement coding Leu.

Table 1 lists the sets of single and double synonyms for those amino acids that can have double synonyms. (We exclude the two synonyms of Ser and the one synonym of Leu for which the reverse complement is Stop, because these do not occur in ambigrammatic genes.)

**Table 1.** For each amino acid (AA) that can have double-synonym single-nucleotide mutations, we list all of the possible codons which do not code for Stop on the complementary strand, indicating their reverse complement (Comp. AA). The codons that have a double synonym are marked with an asterisk. For each of these codons, we list the number of mutations which are synonymous, and the number of double synonym mutations. In each case the numbers of single (double) mutations are written *S*^(n)^ + *S*^(v)^ (*D*^(n)^ + *D*^(v)^), where the superscript n denotes transitions, and superscript v transversions. Also, double synonyms are counted in the list of single synonyms.

| AA | Codon | $S^{(n)} + S^{(v)}$ | $D^{(n)} + D^{(v)}$ | Comp. AA |
| --- | --- | --- | --- | --- |
| Leu | UUG* | 1 + 0 | 1 + 0 | Gln |
|  | CUU | 1 + 1 | 0 + 0 | Lys |
|  | CUC | 1 + 1 | 0 + 0 | Glu |
|  | CUG* | 1 + 2 | 1 + 0 | Gln |
| Pro | CCU* | 1 + 2 | 0 + 1 | Arg |
|  | CCC | 1 + 2 | 0 + 0 | Gly |
|  | CCA | 1 + 2 | 0 + 0 | Trp |
|  | CCG* | 1 + 2 | 0 + 1 | Arg |
| Gln | CAA* | 1 + 0 | 1 + 0 | Leu |
|  | CAG* | 1 + 0 | 1 + 0 | Leu |
| Arg | CGU | 1 + 2 | 0 + 0 | Thr |
|  | CGC | 1 + 2 | 0 + 0 | Ala |
|  | CGA* | 1 + 3 | 0 + 1 | Ser |
|  | CGG* | 1 + 3 | 0 + 1 | Pro |
|  | AGA* | 1 + 1 | 0 + 1 | Ser |
|  | AGG* | 1 + 1 | 0 + 1 | Pro |
| Ser | UCU* | 1 + 1 | 0 + 1 | Arg |
|  | UCC | 1 + 1 | 0 + 0 | Gly |
|  | UCG* | 1 + 2 | 0 + 1 | Arg |
|  | AGU | 1 + 0 | 0 + 0 | Thr |
|  | AGC | 1 + 0 | 0 + 0 | Ala |

#### Implications

Our first rule shows that an ambigrammatic version of any gene can evolve, without making any changes to the amino acid sequence. This establishes how ambigrammatic sequences can arise, but it does not illuminate why they are favoured.

Combined with observed polymorphisms of narnaviruses, the second ambigram rule can give an indication of the utility of ambigrammatic sequences. In studies on the (usual) non-ambigrammatic genomes, the ratio of synonymous to non-synonymous mutations is used as an indicator of whether the nucleotide sequence codes for a protein: non-synonymous mutations are likely to be deleterious if the sequence codes for a functional protein. We shall adapt this approach to our study of ambigrammatic narnavirus genes. We assume that the forward direction is a coding sequence (usually for RdRp), and confine attention to those mutations which are synonymous in the forward direction. If the complementary strand codes for a functional protein, most of these synonymous mutations will inevitably result in changes of the complementary amino acid sequence. However, at many loci the evolutionarily favoured amino acid will be one that allows double synonyms. In these cases, there can be non-deleterious mutations between a pair of codons that preserve the amino acid sequence of both the forward and the complementary strands.

If the complementary strand codes for a functional protein, we expect studies of the polymorphism of the gene would show that these double-synonym loci will be mutational ‘hotspots’, where mutations occur more frequently. In addition, the double-synonym pairs would be represented far more frequently than other mutations at these loci. These observations lead to two distinct tests for whether there is evolutionary pressure on the translated sequence of the complementary strand.

### Ambigrammatic narnavirus genes

We analysed data from samples of two ambigrammatic narnaviruses, *Culex narnavirus 1* (CxNV1, with 46 genomes) and *Zhejiang mosquito virus 3* (ZJMV3, with 10 genomes). Both narnaviruses have an ambigrammatic RdRp coding gene, denoted CxNV1-RdRp and ZJMV3-RdRp respectively. The reverse open reading frame has its codons aligned with the forward frame. In both forward and reverse reading frames any stop codons are close to the 3′ end of the respective frame. The ambigrammatic feature is certainly a puzzle. There appear to be two classes of plausible explanations:

1. **The reverse open reading frame codes a protein**. This is logically possible, but if the RdRp gene is strongly conserved, there is very little flexibility in the rORF. However, in the absence of any additional evidence it is the explanation which requires the fewest additional hypotheses.
2. **The reverse open reading frame facilitates association of ribosomes with RNA**. This could conceivably convey advantages by providing a mechanism to protect viral RNA from degradation, but without further evidence this requires additional hypotheses.

Recently, additional evidence has emerged which may provide support for the second of these explanations. Specifically, the CxNV1 infection has recently been shown to be associated with another ambigrammatic viral RNA segment, termed *Robin* [Batson et al., 2020, Retallack et al., 2021]. It was reported that this segment, CxNV1-Robin, is ambigrammatic, with forward and reverse codons aligned, over very nearly the entire length (about 850 nt), where direction designation is determined by which amino acid sequence appears more conserved. Again, any stop codons occur close to the 3′ end. Neither forward nor reverse directions of Robin are homologous with known sequences.

Because ambigrammatic genes are rare, finding two of them in the same system is a strong indication that their occurrence has a common explanation. This observation makes it appear unlikely that the reverse open reading frame is a device to ‘pack in’ an additional protein coding gene, and more likely that the ambigrammatic feature is associated with allowing ribosomes to associate with both strands of the viral RNA.

This reasoning suggests that the Robin gene may play a role in selecting for the ambigrammatic property (for example, it may facilitate protection by ribosomes of the viral RNA). If this surmise is correct, we should expect to see a version of the Robin gene associated with other ambigrammatic narnaviruses. It is possible that this might be detected by a search of archived sequence data. Only *Zhejiang mosquito virus 3* appeared to be observed multiple times to make detection of an additional Robin segment possible, so we concentrated on that system.

We were able to find evidence of an ambigrammatic RNA, of length approximately 900 nt, that co-occurs with ZJMV3 RdRp segment across multiple samples recovered by at least two studies that, like CxNV1 Robin, bears no recognisable homology (via BLAST [Altschul et al., 1990] or HHpred [Finn et al., 2011]) to publicly available sequences or CxNV1 Robin itself. Given the conjunction of these unusual features we strongly believe this ambigrammatic RNA to be the equivalent of a Robin segment in ZJMV3. We do note, however, that due to contig quality in the datasets where ZJMV3 was found, the typical pentamers found at the ends of narna- [Rodríguez-Cousiño et al., 1998], ourmia- [Wang et al., 2020], and, seemingly, mitovirus [Mizutani et al., 2018] RNA (5′*−*GGGGC and GCCCC*−*3′) cannot be identified unambiguously.

## Methods

### Tests for whether the complementary strand is coding

We have argued that doubly-synonymous mutations will give a signature of the reverse strand coding for a functional protein. If the reverse-direction code is functional, then the only assuredly non-deleterious mutations would be the double-synonym ones, where one codon is transformed by a single-nucleotide substitution to another codon which preserves the amino acid coded in both the forward and the reverse directions.

Assume that we have *M* sequences of an ambigrammatic gene, fully sequenced and maximally aligned with each other, and that one strand, referred to as the ‘forward’ strand, codes for a functional protein. We identify a ‘consensus’ codon at each of the *N* loci, and then enumerate the set of variant codons at each amino acid locus. (The variant set is the set of all codons which were observed at a given locus and which differ from the consensus codon.) If the consensus codon at a locus is one of the twelve double-synonym codons listed in table 1, we term this a *doubly-synonymous locus*. The number of doubly-synonymous loci is *N*_ds_.

There are two different approaches to testing whether double synonyms indicate that the complementary strand is coding:

### Look for the existence of mutational ‘hotspots’

We can look for evidence that the doubly-synonymous loci experience more substitutions than other loci.

For each codon locus *k*, we can determine the number of elements of the variant set, *n*(*k*), (that is, the number of different codons observed at that codon locus which differ from the consensus). We also determine the fraction of codons *f* (*k*) which differ from the consensus codon, that is, the ratio of the number of polymorphs which do not have the consensus codon at site *k* to the total number of polymorphs, *M*). We then determine the averages of these quantities, ⟨*n*(*k*)⟩ and ⟨*f* (*k*)⟩, for the doubly-synonymous loci and for the other loci. If the ratios

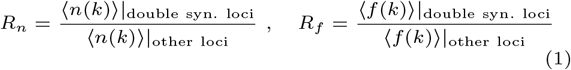

are large, this is evidence that the complementary strand is coding.

The null hypothesis, indicating that the reverse open reading frame is non-coding, is that the ratios *R*_*n*_ and *R*_*f*_ are sufficiently close to unity that the difference may be explained by statistical fluctuations.

### Mutation frequencies test

We can also look at codon frequencies for different mutations at doubly-synonymous loci. If the complementary strand is coding, we expect to find that the frequency of mutations observed at doubly-synonymous loci will heavily favour double-synonym codons over single synonyms. We consider the subset of double-synonym loci where mutations are observed (that is, where *n*(*k*) *>* 1). For each of these *N*_a_ *variable doubly-synonymous loci*, we can determine two numbers: *n*_s_(*k*) is the number of distinct singly-synonymous variants at locus *k*, and *n*_d_(*k*) is the number of these variants which are also doubly-synonymous. (Clearly *n*(*k*) *≥ n*_s_(*k*) *≥ n*_d_(*k*)). If *n*_d_(*k*) = *n*_s_(*k*), that means that the mutations preserve the complementary-strand amino acid, which is an indication that the reverse strand is coding. If {*k*^*^} is the set of variable doubly-synonymous loci, we then calculate

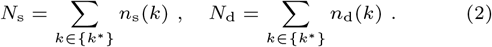

If the complementary strand is coding, we expect

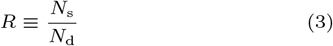

to be close to unity.

However, there will also be beneficial or neutral mutations which do change the amino acids, so that not all mutations will be between sets of doubly-synonymous codons. We need to be able to quantify the extent to which finding other than double-synonym mutations is an indication that the reverse strand is non-coding. We must do this by comparison with a null hypothesis, in which the reverse strand is non-coding.

### Null hypothesis for mutation frequencies

Let *R*_0_ be the value of the ratio *R* that is derived from this null hypothesis that the complementary strand is non-coding. In order to compute the expected *N*_s_*/N*_d_ ratio, *R*_0_, we adopt the following approach. We assume that the *M* sequences are sufficiently similar that only a small fraction of loci have undergone mutations. We adopt the Kimura model [Kimura, 1980], which assumes that the mutation rate *r*_n_ for transitions (A *↔* G or C *↔* U) is different from the rate *r*_v_ for transversions (other single-nucleotide mutations), and negligible for other types of mutation. The ratio of these rates is

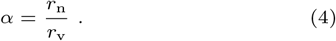

If the numbers of single (double) synonyms of the consensus nucleotide at locus *k* leading to transitions or transversions are respectively 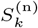 and 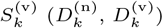, then we estimate

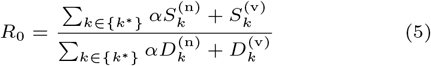

The numbers 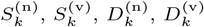 are given in table 1 for all of the double-synonym codons.

### Strandedness of *Culex narnavirus 1* genomes

The Californian mosquito dataset [Batson et al., 2020] from which most *Culex narnavirus 1* sequences came from was prepared using reagents that allow the inference of RNA template strandedness. Based on this we confirmed the expected excess of RNA templates with the same direction as the positive strand (that is, mRNA-sense) of both RdRp and Robin segments of *Culex narnavirus 1* (see Figure 2), since it is a positive-sense single-stranded RNA virus. To ensure this was correct we also checked that some other viruses present in the same dataset also followed expected strand excesses. Two positive sense single-stranded RNA viruses, *Culex flavivirus* and *Marma virus*, and two negative sense single-stranded orthomyxoviruses —*Astopletus* and *Wuhan mosquito virus 6* — did indeed have the expected overwhelming excess of positive-sense reads for the former two, and the moderate excess of negative-sense reads for the latter two [Waldron et al., 2018].

**Fig. 2.**
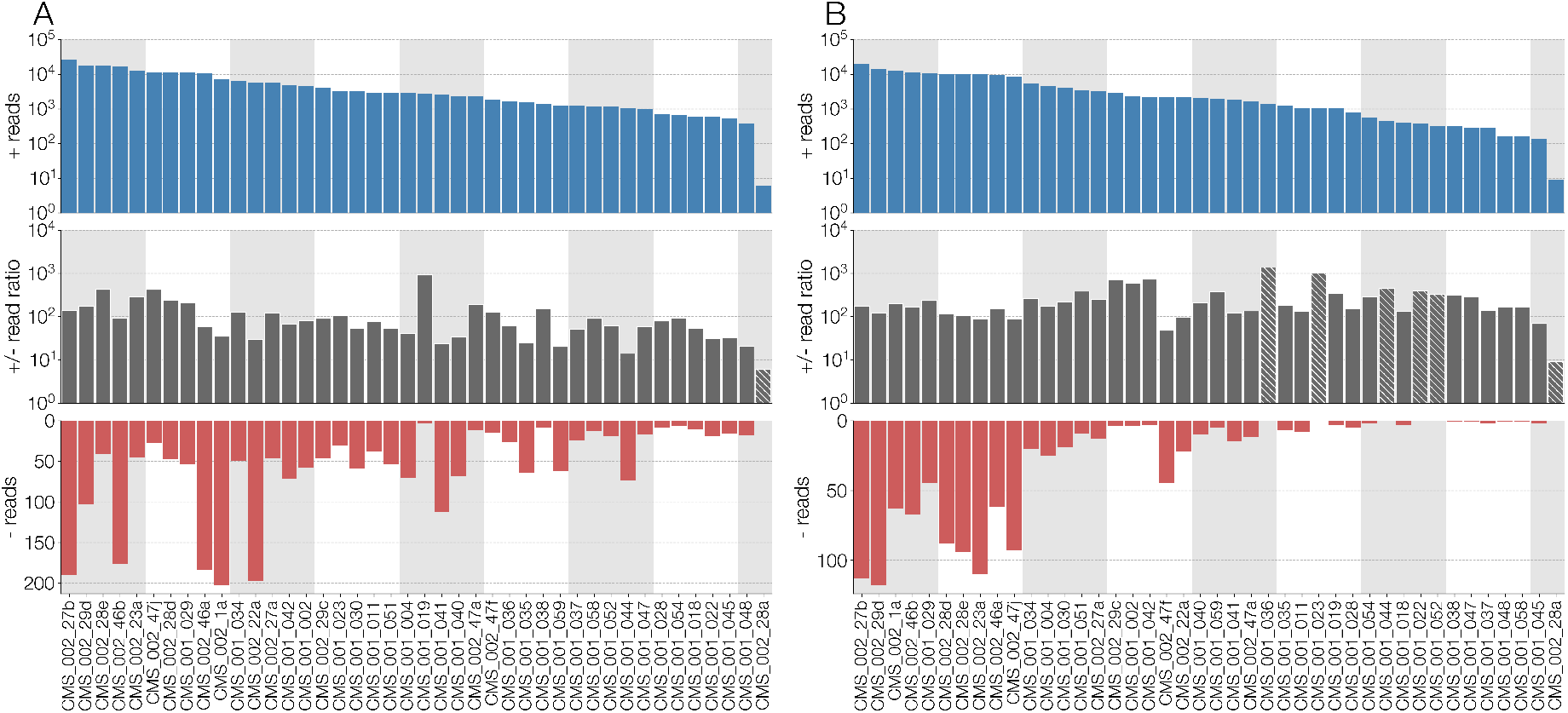
**A**) Number of reads (from top to bottom): in the positive sense (*i*.*e*., same as genomic) with respect to RdRp segment of *Culex narnavirus 1*, the ratio of positive-to negative-sense reads, and negative-sense reads. Numbers of positive-sense reads and ratio of positive-to negative-sense reads are log-scaled while number of negative reads is displayed in normal space. Ratios where no negative sense reads could be identified are highlighted with hatches and the ratio reflects positive-sense reads (*i*.*e*., by assuming there is one negative-sense read). **B**) Same information as **A**, but showing read numbers for Robin segment of *Culex narnavirus 1*. The results are displayed for the 42 different strains, with the samples ordered in descending order of the total number of reads for each segment.

The breakdown of read strandedness was carried out by aligning all non-host reads from each sample in the Californian dataset [Batson et al., 2020] to a consensus sequence of RdRp or Robin segments of *Culex narnavirus 1* with minimap2 [Li, 2018]. Since the library was prepared using reagents that can discriminate the orientation of the original RNA template with respect to a reference, any reverse R1 read (and its forward R2 mate) corresponds to an RNA template in the same direction as the reference, which is positive sense and *vice versa* forward R1 reads (and reverse R2 mate) correspond to a template in the reverse direction (*i*.*e*., negative sense).

### Finding the Robin segment of *Zhejiang mosquito virus 3*

We looked through assembled contig datasets from two metagenomic mosquito studies (three from China and six from Australia) [Shi et al., 2016, 2017], kindly provided to us by Mang Shi and Edward C Holmes. We clustered contigs from the nine datasets by similarity using CD-HIT [Fu et al., 2012] with a threshold of 90% and looked for clusters that contained contigs from at least 6 samples, that did not have standard deviation in contig length greater than 1200, and had fewer than 200 contigs. Of the hundreds of clusters filtered this way only a handful also possessed sequences ambigrammatic across at least 90% of their length and only two clusters were mostly comprised of ambigrammatic sequences, while the rest were clearly recognisable as mosquito contigs. Of the two clusters one was identifiable as the RdRp of *Zhejiang mosquito virus 3*, while we presume the other to be an unrecognisably distant orthologue of *Culex narnavirus 1* Robin, on account of its co-occurrence with ZJMV3 RdRp, ambigrammaticity, and length.

A phylogenetic tree of *Zhejiang mosquito virus 3, Culex narnavirus 1* and their closest relatives was recovered by aligning their RdRp sequences at the amino acid level with MAFFT (E-INS-i setting) [Katoh et al., 2005] and inferring the phylogeny with RAxML [Stamatakis, 2014] under a BLOSUM62+CAT [Henikoff and Henikoff, 1992, Lartillot and Philippe, 2004] amino acid substitution matrix and site rate heterogeneity. The tree (displayed in Figure 1) was rooted with Ochlerotatus-associated narna-like virus 1 sequence, as it appeared to be the longest branch in the tree.

## Results

Next we report the results of our studies of polymorphism of the four ambigrammatic narnavirus genes. We discuss what can be learned from applying standard techniques, before discussing the results of our tests for whether the reverse open reading frame codes for a protein.

### Forward reading frame

Each sequence was trimmed to a length of 3*N* nucleotides. We identified a consensus nucleotide at each locus, and determined the set of variant nucleotides at each locus. We determined the total number of transition and transversion mutations which are observed, *N*_n_ and *N*_v_ respectively. (More precisely, these are the total number of codon loci, across all polymorphs, for which the there is a single-nucleotide mutation relative to the consensus codon which is a transition or transversion.) We also determined the total number of mutations at each position in the codon, (*n*_1_, *n*_2_, *n*_3_). We estimated the average number of variable sites *r* as the total number of nucleotide variants, divided by the product of the number of sequences and alignment length. We also estimated the ratio *α* of the rate of selected transition mutations to the rate of transversions:

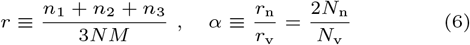

(recall that there are twice as many transversions as transitions). We also determined a ‘normalised’ triplet of variable sites for each position within the codon: (*z*_1_ : *z*_2_ : *z*_3_) = 3(*n*_1_ : *n*_2_ : *n*_3_)*/*(*n*_1_ + *n*_2_ + *n*_3_). Our results on the nucleotide-level investigation of polymorphism are summarised in table 2.

**Table 2.** Nucleotide-level statistics of mutations. The consensus sequence has *N* codons. Among the mutations observed in *M* polymorphs, there are *N*_n_ transitions, *N*_v_ transversions, with overall rate *r* and transition/transversion rate ratio *α*. The numbers total mutations at each base position is (*n*_1_ : *n*_2_ : *n*_3_), and normalising these to ratios via equation (6) yields (*z*_1_ : *z*_2_ : *z*_3_).

| Strand | $N$ | $M$ | $N_n$ | $N_v$ | $r$ | $\alpha$ | $(n_1, n_2, n_3)$ | $(z_1 : z_2 : z_3)$ |
| --- | --- | --- | --- | --- | --- | --- | --- | --- |
| CxNV1-RdRp | 1033 | 46 | 606 | 362 | 0.0068 | 3.35 | (181, 140, 645) | (0.56 : 0.44 : 2.00) |
| ZJMV3-RdRp | 1075 | 12 | 210 | 39 | 0.0064 | 10.80 | (47, 29, 173) | (0.57 : 0.35 : 2.08) |
| CxNV1-Robin | 272 | 46 | 213 | 146 | 0.0096 | 2.92 | (107, 100, 152) | (0.89 : 0.84 : 1.27) |
| ZJMV3-Robin | 304 | 10 | 84 | 48 | 0.0145 | 3.50 | (35, 31, 66) | (0.80 : 0.70 : 1.50) |

We then assigned a consensus codon at each codon locus, selecting the frame by the criterion of minimising the number of stop codons. For each of the *N* codons, we determined the variant set of codons which were observed in each of the *M* sequences. The total number of synonymous and non-synonymous single-nucleotide changes in the variant sets was *N*_sy_ and *N*_ns_ respectively. The total number of mutations (relative to the consensus sequence) encountered in the variant sets where two or three nucleotides were changed was *N*_mult_. For each codon there are numbers of possible non-synonymous mutations which are transistions and transversions, 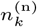 and 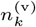, and numbers of synonymous mutations which are transitions and transversions, 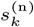 and 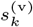 (with 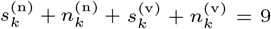). Under the null hypothesis that the sequence is non-coding, the expected value of the ratio

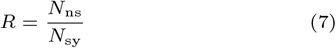

is

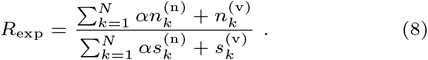

We also determined the fraction of codons where multi-nucleotide mutations are observed, *f*_mult_ = *N*_mult_*/N* . We present our results for the codon-level mutations in table 3, which includes information for both the forward and the complementary read directions (with codon boundaries aligned for the complementary direction).

**Table 3.** Summary of results for codon-level mutations. The numbers of single-nucleotide synonymous and non-synonymous mutations are *N*_sy_ and *N*_ns_ respectively, *N*_mult_ is the number of mutations with more than one base changed, *R*_exp_ is the null value of *R* = *N*_ns_*/R*_sy_, and *f*_mult_ if the fraction of mutations which have multiple-nucleotide changes.

| Strand | $N_{sy}$ | $N_{ns}$ | $N_{mult}$ | $R = N_{ns}/N_{sy}$ | $R_{exp}$ | $R/R_{exp}$ | $f_{mult}$ |
| --- | --- | --- | --- | --- | --- | --- | --- |
| CxNV1-RdRp-fwd | 623 | 189 | 123 | 0.303 | 2.37 | 0.128 | 0.12 |
| ZJMV3-RdRp-fwd | 170 | 59 | 13 | 0.347 | 2.14 | 0.162 | 0.012 |
| CxNV1-Robin-fwd | 112 | 141 | 89 | 1.26 | 2.34 | 0.538 | 0.45 |
| ZJMV3-Robin-fwd | 49 | 61 | 14 | 1.24 | 2.35 | 0.528 | 0.046 |
| CxNV1-RdRp-comp | 136 | 676 | 123 | 4.97 | 2.43 | 2.04 | 0.12 |
| ZJMV3-RdRp-comp | 50 | 179 | 13 | 3.58 | 2.14 | 1.67 | 0.012 |
| CxNV1-Robin-comp | 66 | 187 | 89 | 2.83 | 2.39 | 1.23 | 0.45 |
| ZJMV3-Robin-comp | 32 | 78 | 14 | 2.43 | 2.28 | 1.07 | 0.046 |

The alignments are *ambigrammatic*, in the sense that there are no stop codons in the interior of the sequence. None of the individual sequences had stop codons in the body of the sequence in either direction.

We also computed ORF-wide d*N* /d*S* values (plotted in figure 1(**b**)), by assuming that every mutation in the alignment has occurred only once to be conservative. This was motivated by the presence of pairs of sites with four haplotypes between them (4G sites), an indication that recombination may be a potential issue with narnavirus sequences. Normalising the number of observed non-synonymous and synonymous mutations was done by assuming a transition/transversion ratio of 2, consistent with equation (6). These values d*N* /d*S* values are slightly different from the *R/R*_exp_ ratios in table 3 because the latter excludes mutations where more than one base differs from the consensus codon. In all but one of the cases d*N* /d*S* is higher than *R/R*_exp_, because the multiple nucleotide mutations which are included in d*N* /d*S* are predominantly non-synoymous.

Based upon these tables, we can make the following observations and deductions:

1. **Diversity**. We observe that both RdRp and Robin segments are comparable in their diversity, for both CxNV1 and ZJMV3. As expected, RdRp sequences are highly conserved at the amino acid level. Robin, on the other hand, appears far more relaxed at the amino acid level and, consistent with this, diverged beyond recognition between CxNV1 and ZJMV3.
2. **Relative mutation rate by codon position**. For RdRp sequences, more mutations are observed at the third nucleotide in each codon, as expected for a sequence that preserves the amino acid sequence (because most synonymous mutations involve the third nucleotide of a codon). In the case of Robin sequences, the frequencies of mutation are much closer to being equal, to the extent that for CxNV1-Robin the null hypothesis that the rates are equal is not definitively rejected. However, mutations at different codon sites are sufficiently weighted towards the third position that we shall assume that Robin does code for a functional protein. While the values of (*z*_1_ : *z*_2_ : *z*_3_) are very different for RdRp and Robin, their values are comparable for CxNV1 and ZJMV3, which is an indication that the selective pressures on both viruses are the same.
3. **Rate of multiple-nucleotide mutations**. The fraction of multiple-nucleotide mutations is higher for Robin sequences than it is for RdRp sequences. This may be an indication that the Robin sequence is under strong selective pressure, because some aminoacid substitutions can only be achieved through multiple nucleotide mutations.
4. **Transition to transversion ratio**. Three of the values of *α* were similar to each other, while the value for ZJMV3-RdRp was higher than the others. Because transitions occur at a higher intrinsic rate, a lower value of *α* indicates that observed mutations are biased in favour of the rarer transversions, which is an indication of unusual selective pressures. The fact that the values of *α* for the Robin segments are comparable to, or lower than, the values for RdRp are a further indication that Robin is under similar selective pressure too.
5. **Ratio of non-synonyms to synonyms**. For the RdRp segments the values of *R* = *N*_ns_*/N*_s_ are much smaller than the values *R*_0_ predicted (equation (8)) by the null hypothesis that mutations are random. This indicates that the selective pressure on RdRp acts to preserve the amino acid sequence. For Robin segments, the values of *R* are much larger, but still smaller than the prediction from the null hypothesis. This indicates that while points 1-4 above indicate that Robin is under some selective pressure, the amino acid sequence is not strongly conserved. This is consistent with the hypothesis that the selection acting on Robin is relaxed. Figure 3 illustrates the distribution of mutations across the forward and reverse reading frames of all four ORFs for both CxNV1 and ZJMV3. As expected, there is evidence that some regions accumulate mutations more readily than others. The pattern is consistent with what would be expected from the statistical reductions in the tables.

**Fig. 3.**
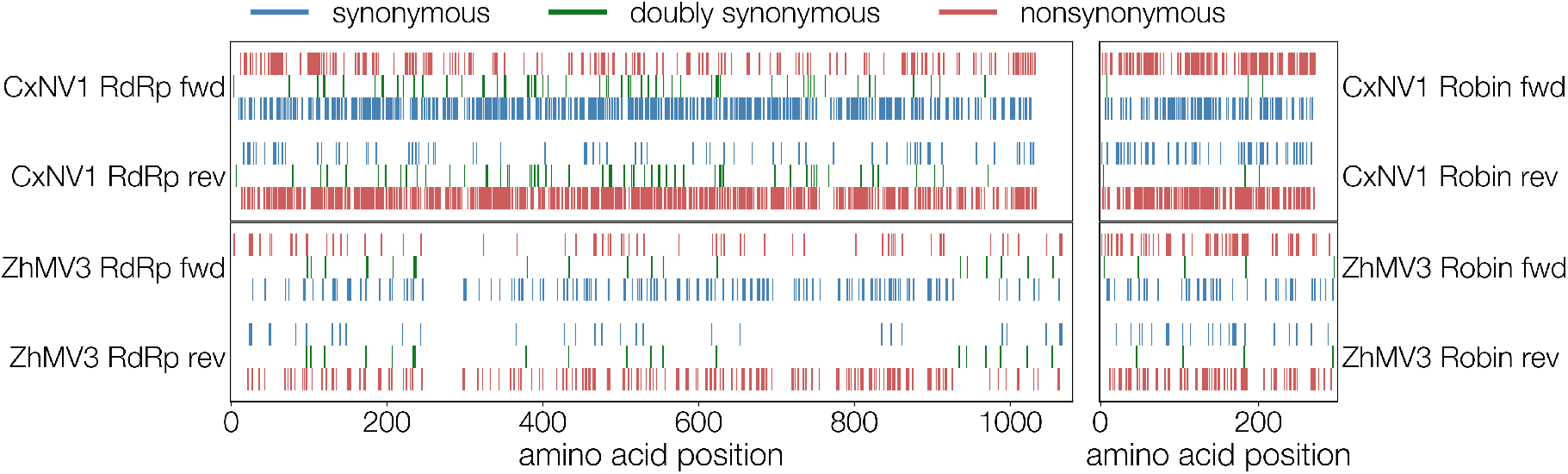
Distribution of synonymous (blue), non-synonymous (red) substitutions, and doubly synonymous sites (green) in CxNV1 (upper plots) and ZJMV3 (lower plots) RdRp (left) and Robin (right) segments in both directions (forward towards top, reverse towards bottom). Translated reverse ORFs are shown backwards (segment coordinate space). Double synonyms don’t overlap perfectly because forward and reverse ORFs differ in length and begin and end at different positions along the segment.

### Complementary reading frame

We determined the set of *N*_ds_ doubly-synonymous codons in the consensus sequence, and the subset of *N*_a_ of these which have variant codons.

1. **Mutational hotspots test**. We applied the mutational hotspots test to all four sequences, as described by equations (1) above. The results (tables 4) show no evidence that the doubly synonymous sites are undergoing more frequent mutations, or that their mutations are more widely spread across the dataset.
2. **Mutation rate test**. We examined the number of mutations in the set of *N*_a_ doubly-synonymous sites which were variable. We found (table 5) that many more of the observed mutations at these sites are only singly synonymous, when a doubly-synonymous mutation is possible, which is further evidence that the complementary strand is non-coding. The numbers of doubly-synonymous mutations were quite low, and so it was not possible to make a reliable comparison of the ratio *N*_s_*/N*_d_ with the null hypothesis.
3. **Ratio of non-synonyms to synonyms** The ratios of non-synonymous to synonymous mutations, presented in table 3 and figure 1(**b**), were lower than the null hypothesis for the forward direction. This is readily explained as an indication that the forward ORF codes for a functional protein. However the *N*_ns_*/N*_s_ ratios for the reverse direction were all higher than the null hypothesis. This observation is explained, qualitatively, as follows. If the forward direction strictly conserves the amino acid sequence, then all of the mutations which are synomymous on the reverse strand are doubly-synonymous. Because only 12 of the 64 codons allow for doubly-synonymous mutations, the *N*_ns_*/N*_s_ ratio would be very high for the complementary strand if the forward sequence were to be exactly conserved. We computed this ratio, and found 11.2 for CxNV1-RdRp, and similar values for the other sequences. This theoretical ratio is considerably higher than the measured value of 4.97, because the forward sequence is not exactly conserved. For Robin segments, the value of *R* for the reverse ORF is only slightly higher than the null hypothesis, because the amino acid sequence is only weakly conserved.

**Table 4.** Summary of results of the mutational hotspots test. Left panel: values of the average number of elements of the variant set, ⟨*n*(*k*⟩ and of the average fraction of non-consensus codons, ⟨ *f* (*k*)⟩, for double-synonym sites, and for the other sites. Right panel: *N* is the number of loci in the alignment, *N*_ds_ is the number of double-synonym loci, and *R*_*n*_, *R*_*f*_ are the ratios of ⟨*n*(*k*) ⟩ and ⟨*f* (*k*) ⟩ at double-synonym sites to their values at other sites. The differences of these ratios from unity do not appear significant.

| Sample | $\langle n(k) \rangle$ | $\langle f(k) \rangle$ |
| --- | --- | --- |
| Double syns., CxNV1-RdRp | 0.954 | 0.161 |
| Other codons, CxNV1-RdRp | 0.968 | 0.155 |
| Double syns., ZJMV3-RdRp | 1.20 | 0.042 |
| Other codons, ZJMV3-RdRp | 1.23 | 0.050 |
| Double syns, CxNV1-Robin | 1.76 | 0.195 |
| Other codons, CxNV1-Robin | 1.48 | 0.169 |
| Double syns, ZJMV3-Robin | 0.889 | 0.096 |
| Other codons, ZJMV3-Robin | 0.960 | 0.097 |

**Table 5.** Results for the mutational codon frequency test: *N* is the number of loci in the alignment, *N*_a_ is the number of mutationally active double-synonym loci, and *N*_s_, *N*_d_ are, respectively, the numbers of single and double synonym mutations. The actual ratio *R* = *N*_s_*/N*_d_ is compared with the null-hypothesis value *R*_0_, equation (8).

| Sample | $N$ | $N_a$ | $N_s$ | $N_d$ | $R$ | $R_0$ | $R/R_0$ |
| --- | --- | --- | --- | --- | --- | --- | --- |
| CxNV1-RdRp | 1033 | 136 | 151 | 60 | 2.51 | 3.02 | 0.83 |
| ZJMV3-RdRp | 1075 | 219 | 33 | 20 | 1.65 | 3.21 | 0.51 |
| CxNV1-Robin | 272 | 40 | 24 | 3 | 8.00 | 3.21 | 2.49 |
| ZJMV3-Robin | 304 | 59 | 20 | 4 | 4.00 | 4.04 | 0.99 |

## Discussion

We have argued that doubly synonymous codons provide a key to understanding whether ambigrammatic viral RNA segments code for two functional proteins. If there were two coding genes, doubly synonymous mutations would be mutational hotspots, because they are unambiguously non-deleterious. We applied our analysis to recent observations of polymorphisms in two ambigrammatic narnaviruses: *Culex narnavirus 1* and *Zhejiang mosquito virus 3*. There was no evidence that doubly synonymous sites are mutational hotspots, or that there is a prevalence of mutations to other doubly-synonymous codons at these sites. Other, circumstantial, evidence favours the interpretation that the complementary strand is non-coding. Ambigrammatic sequences have been observed in other narnaviruses, but they are undoubtedly a rare phenomenon. If the rORF (reverse open reading frame) of both RdRp and Robin segments had evolved to code for a functional protein, each RNA segment would code for two genes. Given that ambigrammatic sequences are rare [DeRisi et al., 2019], finding a system where two had evolved independently would be highly improbable. Moreover, because the ambigrams are full length, each of the ambigrammatically coded sequences would code for two genes which have the same length as each other.

An observation of the simultaneous detection of two or more ambigrammatic genes would strongly favour models where there is an advantage in evolving an ambigrammatic sequence which is independent of whether the reverse open reading frames are translated into functional proteins. This argument led us to discover the Robin segment of ZJMV3, and suggests that more ambigrammatic narnaviruses with at least two segments will be discovered by metagenomic surveys, when suitable data sets become available. Similarly, the elusive Robin segment should already be hiding in datasets of narnaviruses descended from the common ancestor of CxNV1 and ZJMV3.

Our studies of polymorphisms in the forward direction indicate that both RdRp and Robin are under purifying selection. In the case of RdRp the amino acid sequence is strongly conserved, but the Robin sequence is not.

The role of the RdRp coding fragment is already understood. This makes it plausible that the other fragment plays a role which facilitates the evolution of ambigrams. If the lack of stop codons on the complementary strand is not required to allow protein synthesis, we can surmise that its role is to allow ribosomes to associate with the complementary strand. Having RNA segments able to be covered by ribosomes may provide some protection for the viral RNA against degradation.

Recent experiments indicate that ambigrammatic narnavirus genes display unusual ribosome profiles, with a ‘plateau’ structure [Retallack et al., 2021]. It has been argued [Wilkinson et al., 2021] that the plateaus indicate that the ribosomes attached to the viral RNA become stalled, creating a cover (see also Cepelewicz [2020]). The ambigram property allows binding of ribosomes to both strands, hiding the viral RNA from host defence and degradation mechanisms. We can surmise that there exists a molecule which binds to the 3′ end of the viral RNA, preventing release of ribosomes [Wilkinson et al., 2021]. It is possible that Robin plays a role in this process, by creating a protein which blocks ribosome detachment at 3′ end. Alternatively, it might be proposed that the ribosome ‘traffic jam’ is a consequence of the structure of the RdRp itself, due to formation of RNA hairpins. However, these would have to trade off against RdRp function. The proposed mechanism involving Robin making a blocking protein has the advantage that the RdRp works efficiently when the viral RNA concentration is small. Later, after it has duplicated many copies of itself and of Robin, the Robin protein attaches to the viral RNA and creates stalled polysomes, protecting the viral RNA from degradation.

| Gene | $N$ | $N_{ds}$ | $R_n$ | $R_f$ |
| --- | --- | --- | --- | --- |
| CxNV1-RdRp | 1033 | 220 | 0.986 | 1.044 |
| ZJMV3-RdRp | 1075 | 219 | 0.975 | 0.840 |
| CxNV1-Robin | 272 | 54 | 1.19 | 1.16 |
| ZJMV3-Robin | 304 | 81 | 0.926 | 0.978 |

There may, however, be additional viral genes involved in ambigrammatic narnavirus infections, and there are many possible roles for the Robin gene. It could code a protein which inhibits the mechanism of ‘no-go-decay’, which releases stalled ribosomes, play a role in the viral suppression of RNAi [Mierlo et al., 2014] or in formation of syncytia or viral particles. Without a better understanding of the narnavirus lifecycle in arthropods it is not certain whether Robin does code for a protein which blocks detachment of ribosomes.

We did search the CxNV1 dataset for further fragments of ambigrammatic viral RNA, which might be candidates for coding additional genes. A search for additional ambigrammatic sequences greater than 200nt in length did not produce any candidates.

A recent preprint [Retallack et al., 2021] presents evidence that peptides translated from the reverse ORF of the RdRp segment can be detected (though not quantified) and that inserting mutations in the RdRp sequence which are synonymous in the forward reading frame but introduce stop codons in the reverse frame reduces the fitness of the virus. The mutations were clustered close to the 3′ end of the RdRp gene. These observations could be interpreted as indicating that the reverse reading frame codes for a functional protein or that all ORFs in the cell may be translated in a ‘leaky’ way. However, changing the RNA sequence may also interfere with the action of molecules which bind to the RdRp strand.

The data on strandedness indicates that there are mechanisms which favour the presence of one strand rather than the other. Our results do not suggest that this is related to the ambigrammatic property, and are consistent with the strandedness bias being determined by mechanisms which are also found in other virus families. Narnaviruses are distant relatives of the RNA bacteriophage family *Leviviridae* and of eukaryote-infecting mitoviruses that replicate in mitochondria, while ourmiaviruses are most closely related [Shi et al., 2016]. It has been suggested that narnaviruses [Rodríguez-Cousiño et al., 1998], as well as their relatives ourmia-, mito-, and leviviruses, possess conserved secondary RNA structures at their ends. in narnaviruses these structures, particularly at the 3′ end, have been shown to proect genomic RNA from exonuclease degradation [Esteban et al., 2005]. We may suppose that these secondary RNA structures could also function in regulating translation, though understandably the corpus of narnavirus molecular biology research is scarce. It may be tempting to rely on similar genomic features discovered in distant relatives of narnaviruses, namely RNA bacteriophages in the family *Leviviridae*, that have been investigated in far greater detail, with the caveat that these viruses possess a markedly different genomic organisation and infect entirely different hosts, and thus may prove more useful conceptually.

## Author contributions

GD devised and directed the search for an analog of Robin in the ZJMV3 sequence archive. MW produced a draft of the manuscript following discussions with the other authors about the recent discovery of a narnavirus system which has two ambigrammatic genes. All authors contributed to writing the manuscript, and reviewed the manuscript before submission.

## Acknowledgments

We thank Hanna Retallack and Joe DeRisi for discussions of their experimental studies of narnaviruses and Amy Kistler for assistance with narnaviral genomes. We thank Amy Kistler and John Pak for comments on a draft. We would like to thank Mang Shi and Edward C Holmes for sharing assembled contigs from Australian and Chinese mosquito metagenomic datasets. G.H. and D.Y. were supported by the Chan Zuckerberg Biohub; MW thanks the Chan Zuckerberg Biohub for its hospitality.

## Data availability

